# Widespread latent hyperactivity of nociceptors outlasts enhanced avoidance behavior following incision injury

**DOI:** 10.1101/2024.01.30.578108

**Authors:** Alexis G. Bavencoffe, Elia R. Lopez, Kayla N. Johnson, Jinbin Tian, Falih M. Gorgun, Breanna Q. Shen, Michael X. Zhu, Carmen W. Dessauer, Edgar T. Walters

## Abstract

Nociceptors with somata in dorsal root ganglia (DRGs) exhibit an unusual readiness to switch from an electrically silent state to a hyperactive state of tonic, nonaccommodating, low-frequency, irregular discharge of action potentials (APs). Ongoing activity (OA) during this state is present in vivo in rats months after spinal cord injury (SCI), and has been causally linked to SCI pain. OA induced by various neuropathic conditions in rats, mice, and humans is retained in nociceptor somata after dissociation and culturing, providing a powerful tool for investigating its mechanisms and functions. An important question is whether similar nociceptor OA is induced by painful conditions other than neuropathy. The present study shows that probable nociceptors dissociated from DRGs of rats subjected to postsurgical pain (induced by plantar incision) exhibit OA. The OA was most apparent when the soma was artificially depolarized to a level within the normal range of membrane potentials where large, transient depolarizing spontaneous fluctuations (DSFs) can approach AP threshold. This latent hyperactivity persisted for at least 3 weeks, whereas behavioral indicators of affective pain – hindpaw guarding and increased avoidance of a noxious substrate in an operant conflict test – persisted for 1 week or less. An unexpected discovery was latent OA in neurons from thoracic DRGs that innervate dermatomes distant from the injured tissue. The most consistent electrophysiological alteration associated with OA was enhancement of DSFs. Potential in vivo functions of widespread, low-frequency nociceptor OA consistent with these and other findings are to amplify hyperalgesic priming and to drive anxiety-related hypervigilance.

## 1. Introduction

Pain after bodily injury is often accompanied by complex behavioral and neural alterations that are poorly understood ^12, 58^, and may cause clinical problems long after surgery or trauma ^28, 37,59^. Ongoing behavioral effects manifested acutely after significant tissue injury include spontaneous grimacing, guarding behavior, and reduced locomotor activity ^12, 60, 67, 73^, while a prominent ongoing neural effect is spontaneous electrical activity in primary afferent neurons ^10, 25, 39, 62, 75^. Many painful effects require extrinsic stimulation to reveal altered sensation and behavior, including allodynia during innocuous stimulation and hyperalgesia during noxious stimulation. Although injury that causes nerve damage can induce chronic neuropathic pain ^27^, injuries that spare larger nerve branches produce ongoing pain, hyperalgesia, and allodynia that often resolve within a week or two ^60^. Nevertheless, latent effects may remain for additional weeks or longer, as illustrated by persistent hyperalgesic priming (revealed during subsequent inflammatory challenge) ^36, 64^ and by persistent latent sensitization of nocifensive reflexes masked by constitutive activity of endogenous analgesic systems ^20, 48^.

Although both hyperalgesic priming and latent sensitization are associated with alterations within the central nervous system (CNS) ^19, 20, 36, 57, 65^, each has also been linked to alterations in primary nociceptors ^30, 38, 50, 64, 66^. Indeed, allodynia and hyperalgesia in many painful conditions depend upon central sensitization induced and/or maintained by ongoing primary afferent activity ^7, 13, 32, 49^. For injuries such as incision in which hyperalgesic priming and latent sensitization outlast ongoing pain, a logical possibility is that persistent readiness to re-enter a painful sensitized state might be promoted by hyperexcitable primary nociceptors poised to generate sustained ongoing activity (OA). Transition from latent hyperactivity to tonic OA in a sufficient number of nociceptors for a sufficient period could redrive central sensitization to promote the allodynia, hyperalgesia, and ongoing pain that sometimes return after healing and resolution of pain ^28, 64^. Consistent with this possibility is enhancement of hyperexcitability (indicated by decreased action potential threshold) reported in nociceptors during in vitro application of PGE_2_ 8 days after in vivo induction of hyperalgesic priming by fentanyl, although this effect was attributed to altered PGE_2_ signaling in nociceptors rather than an alteration of intrinsic excitability ^38^. For tissue injury, evidence is lacking for electrophysiological alterations in nociceptors that persist in latent form after the resolution of ongoing pain and behavioral sensitization, and especially for alterations that enable a transition from electrical silence to sustained OA (hyperactivity).

Making use of the unusual and experimentally useful capacity of nociceptor somata to retain hyperexcitable alterations induced in vivo after subsequent cellular dissociation and culturing ^10, 42, 43, 45, 52, 68, 70, 72^, we show that a latent hyperactive state persists in widespread nociceptors for weeks after unilateral or bilateral hindpaw incision injury, outlasting ongoing mechanical allodynia and hyperalgesia that are evident in voluntary avoidance behavior. Several unexpected features of latent hyperactivity after injury point to nociceptor functions that go beyond ongoing stimulation of affective pain.

## 2. Methods

### 2.1. Animals

All procedures followed the guidelines of the International Association for the Study of Pain and were approved by the McGovern Medical School Animal Care and Use Committees. Male Sprague-Dawley rats (Envigo, USA), 8-9 weeks old, 250-300g, housed 2 per cage were used for the entire study. We elected to postpone investigation of potential sex differences to reduce the number of factors that could interact with many other potential factors in vivo and in vitro that might impact the expression of electrophysiological effects of plantar incision. In our initial sets of experiments, all on unilateral plantar incision (Sections 4.1, 4.2, 4.3), animals acclimated to a 12-hour reversed light/dark cycle for at least four days before beginning experiments, and the rats received all manipulations and tests in their dark active phase. In the remaining studies, the rats were on a normal light/dark cycle, and all manipulations and tests occurred in the rats’ inactive light phase.

### 2.2. Surgical procedures

Animals were randomly assigned to surgical and naïve control groups. Previous studies of plantar incision have shown no effects of sham treatment (reproducing all procedures but omitting incision) on behavioral, physiological, or cellular measures. Unilateral plantar incision was performed using a standard procedure ^14, 15, 82^. Briefly, anesthesia was induced by placing the rat in a chamber filled with 4% isoflurane in air followed by 2% isoflurane via a nose cone maintained during surgery. The left hind paw was sterilized with alternating iodine swabs and alcohol pads. Beginning 0.5-1 cm from the proximal edge of the heel, a longitudinal incision was made through the skin and fascia with a no. 10 scalpel blade. The flexor digitorum brevis muscle was then elevated and incised longitudinally (∼1 cm), leaving intact the muscle origin and insertion. The skin was closed with two absorbable sutures.

A novel bilateral extended plantar incision (EPI) model included the same longitudinal incision through skin, fascia, and flexor digitorum brevis muscle as in the unilateral plantar incision model, but this was given to both hindpaws. Each plantar injury was then extended modestly by making a small incision transversely across the widest part of each of the two most distal pads (the second and third interdigital volar pads, see schematic diagrams in **Figures 1-6**) through the thick pad epidermis and the underlying adipose tissue without cutting into underlying tendons and bone. The pad incisions were not sutured, but hemostasis was promoted by gentle pressure on the wound.

**Figure 1.**
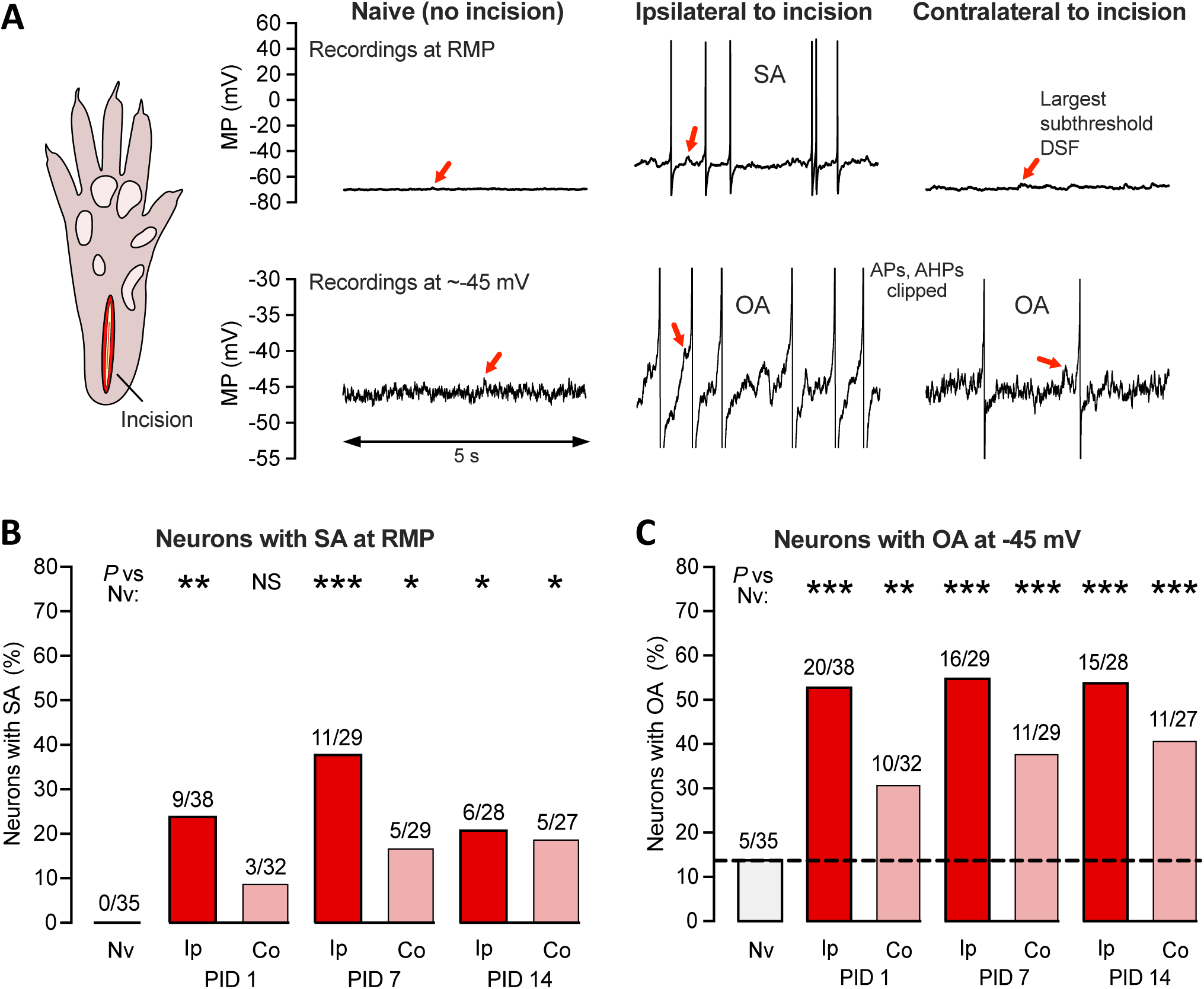
Unilateral plantar incision induces long-lasting, widespread hyperexcitability that can be observed after dissociation of DRG neurons. (A) Representative 5-second traces from somal recordings of neurons isolated from DRGs of naïve rats or DRGs ipsilateral or contralateral to a unilateral plantar incision (schematic on left) made 7 days before dissociation. Recordings in the upper panels were made at RMP and show the entire APs, while those in the lower panels were made at a holding potential of approximately −45 mV, where large DSFs can sometimes reach AP threshold, and are enlarged (clipping APs and afterhyperpolarizations) to be better show individual DSFs. The largest subthreshold DSF in each trace is indicated by a red arrow. (B) Proportion of neurons exhibiting SA (OA at RMP) when dissociated after the indicated number of postinjury days (PID) following unilateral deep plantar incision. Above each bar are the number of neurons exhibiting SA/the number of neurons sampled for that group. Fisher’s exact tests versus naïve group were performed at 3 time points, and with the significance levels Bonferroni-corrected for multiple comparisons. (C) Proportion of the same neurons as in panel B that exhibited OA at −45 mV. AHP, afterhyperpolarization; AP, action potential; Co, contralateral to injury; DSF, depolarizing spontaneous fluctuation; Ip, ipsilateral; Nv, naïve; MP, membrane potential; NS, not significant; OA, ongoing activity; PID, postinjury day; SA, spontaneous activity.

Following surgery, the rat remained on an insulated heating pad until full recovery from anesthesia before being moved back to its cage and observed periodically until full ambulatory recovery. All wounds stopped bleeding within ∼5 minutes and appeared outwardly to have largely healed within 5-6 days, similar to what has been described for unilateral plantar incision ^15, 31^. Simple schematic diagrams are included in each results figure to show the type of incision injury used.

### 2.3. Dissociation and culture of DRG neurons

A dissociated DRG neuron preparation was used that we have repeatedly found in nociceptors from rats, mice, and humans to retain injury-induced, pain-related hyperexcitability for more than 24 hours in culture after in vivo induction of persistent neuropathic pain ^8–11, 42, 52, 53, 81, 84^. Rats were euthanized using intraperitoneal injection of pentobarbital/phenytoin solution (Euthasol, 0.9 ml, Virbac AH, Inc.) followed by transcardial perfusion of ice-cold phosphate buffered saline (PBS, Sigma-Aldrich). DRGs were harvested from below the vertebral T10 level to L6. Ganglia were surgically desheathed before transfer to high-glucose DMEM culture medium (Sigma-Aldrich) containing trypsin TRL (0.3 mg/ml, Worthington Biochemical Corporation) and collagenase D (1.4 mg/ml, Roche Life Science). After 40-min incubation under constant shaking at 34°C, digested DRGs were washed by two successive centrifugations and triturated with two fire-polished glass Pasteur pipettes of decreasing diameters. Cells were plated on 8-mm glass coverslips (Warner Instruments) coated with poly-L-ornithine (Sigma-Aldrich) 0.01% in DMEM without serum or growth factors, and incubated overnight at 37°C, 5% CO2 and 95% humidity.

### 2.4. Recording from dissociated DRG neurons

Recordings were made from isolated neurons with soma diameters ≤ 30 µm, which under our conditions are capsaicin-sensitive in ∼70% and bind isolectin B4 in >30% of sampled neurons ^10, 53, 81^. We did not sample larger neurons because our major goal was to define mechanisms important for SA in probable nociceptors that have been linked to ongoing neuropathic pain, and in vivo recordings after SCI in rats showed that chronic, low-frequency SA was only generated in the somata of C-fiber and Aδ-fiber DRG neurons, which have relatively small soma diameters^10^. Whole-cell patch-clamp recordings were performed at room temperature (∼ 21°C) 18-30 hours after dissociation using an EPC10 USB (HEKA Elektronik) amplifier and a Multiclamp 700B (Molecular Devices). Neurons were observed at 20x magnification on an IX-71 (Olympus, Japan) inverted microscope and recorded in an extracellular solution (ECS) containing (in mM) 140 NaCl, 3 KCl, 1.8 CaCl_2_, 2 MgCl_2_, 10 HEPES, and 10 glucose, which was adjusted to pH 7.4 with NaOH and to 320 mOsM with sucrose. Patch pipettes were made of borosilicate glass capillaries (Sutter Instrument) with a horizontal P-97 puller (Sutter Instrument) and fire polished with a MF-830 microforge (Narishige) to a final pipette resistance of 3-8 MΩ when filled with an intracellular solution composed of (in mM) 134 KCl, 1.6 MgCl_2_, 13.2 NaCl, 3 EGTA, 9 HEPES, 4 Mg-ATP, and 0.3 Na-GTP, which was adjusted to pH 7.2 with KOH and 300 mOsM with sucrose. After obtaining a tight seal (>3 GΩ), the plasma membrane was ruptured to achieve whole-cell configuration under voltage clamp at −60 mV. Recordings were acquired with PatchMaster v2x90.1 (HEKA Elektronik) and Clampex 10.7 (Molecular Devices). The liquid junction potential was calculated to be ∼4.3 mV and this estimate was not corrected, meaning the actual potentials were up to ∼4.3 mV more negative than indicated in the recordings and reported membrane potentials. Identification of nonaccommodating (NA) and rapidly accommodating (RA) neurons was done by stimulating primary neurons with an ascending series of 2-second pulses (increment 5-20 pA) to two times rheobase ^53^. If repetitive discharge was observed, the neuron was classified as NA. If only a single AP was observed (at the beginning of the step), the neuron was classified as RA. RA neurons, which appear to lack the capacity for OA ^53^ and always represented a minority of sampled neurons, were excluded from further analysis. AP voltage threshold and rheobase (current threshold) were also determined during the 2-second depolarizing step protocol. OA at RMP (intrinsically generated spontaneous activity [SA]) was defined as any discharge occurring during the 60-second current clamp recording (I = 0) after first switching to current clamp mode ^10^. OA at −45 mV was defined as any discharge occurring during a 30-second current clamp recording with the neuron artificially held at −45 mV. This test was performed 1-2 minutes after the 2-second depolarizing step protocol ^53^.

### 2.5. Quantifying depolarizing spontaneous fluctuations of membrane potential (DSFs)

Whole-cell current-clamp recordings were analyzed using the Frequency-Independent Biological Signal Identification (FIBSI) program ^17^. FIBSI was written using the Anaconda (v2019.7.0.0, Anaconda, Inc) distribution of Python (v3.5.2) with dependencies on the NumPy and matplotlib.pyplot libraries. It incorporates the previously published algorithm used to analyze DSFs in our prior publications ^11, 29, 42, 47, 53^. FIBSI uses the Ramer-Douglas-Peucker algorithm to detect DSFs, which were obtained from 30-s recordings sampled at 10 kHz with PatchMaster v2x90.1 (HEKA Elektronik) or Clampex 10.7 and filtered with a 10 kHz Bessel filter. A user-defined 1-s sliding median function built into FIBSI was used to estimate resting membrane potential (RMP) at each coordinate, and then FIBSI returned the coordinates, amplitudes, and durations of identified APs and DSFs and hyperpolarizing spontaneous fluctuations (HSFs) (each type with minimum amplitude and duration of 1.5 mV and 5 ms, respectively). Amplitudes for the suprathreshold DSFs eliciting APs were estimated conservatively as the difference between the most depolarized potential reached by the largest subthreshold DSF within the recording and the sliding median at that point. A minimal interval of 200 ms between any two APs was required for the second AP to be substituted by the maximal suprathreshold DSF amplitude in the recording period. If the interval between two APs was less than 200 msec, then a substitution was performed only if a clear peak of a separate DSF or HSF was detected between the APs ^53^. The FIBSI source code and detailed tutorial are freely available for use, modification, and distribution on a Github (GitHub, Inc.) repository titled “FIBSI Project” by user “rmcassidy” (https://github.com/rmcassidy/FIBSI_program).

### 2.6. Behavioral testing

#### Measurement of guarding behavior

Guarding was assessed using methods adapted from previous reports ^15, 83^. Each rat was placed on an elevated wire mesh surface (6 x 6 mm grid size) and confined by a clear plastic cage top (20.5 x 20.5 x 13.5 cm). Both hind paws were assessed by a blinded observer during twelve 1-minute periods every 3 to 4 minutes for 36 to 48 minutes, which were conducted 1, 7, and in some cases 14 days after either deep plantar incision, skin incision, or no treatment (naïve group). The position and condition of each hind paw maintained for at least 30 seconds in each 1-minute observation period were scored on a combined 3-point scale. A score of 0 was given if the incised area was touching and blanched or distorted by the mesh, 1 if the incised area was touching the mesh without blanching or distortion, and 2 if the incised area was elevated entirely off the mesh. For each hind paw the 12 scores were summed and the normalized guarding score was obtained for each rat by subtracting the summed score of the uninjured paw from that of the contralateral injured paw.

#### Operant mechanical conflict (MC) test

The 3-chambered Mechanical-Conflict System (MCS, Coy Laboratory Products; now sold by Noldus, Wageningen, the Netherlands) was used for operant assessment of aversion to noxious mechanical stimulation of the paws ^33, 54, 56^. Unlike other studies, which measure the latency of extensively habituated rats to cross noxious probes a single time, our major measure of altered pain sensation was the number of times a rat voluntarily crossed a chamber containing a dense array of sharp probes elevated to either 4 mm (previously shown to be noxious, providing a measure of mechanical hyperalgesia), 1 mm (innocuous, providing a measure of mechanical allodynia), or 0 mm (absent) ^54^. Furthermore, we departed from our earlier study and other studies using the MCS by omitting the light that triggers photophobia, which had been assumed to be the major motivator for crossing the sharp probes. Instead, our MC test relies solely on the conflict between aversive pain evoked by stepping on noxious probes and a rat’s strong drive to exhaustively explore a relatively unfamiliar environment. On Day 1 rats were acclimated to the dark testing room (illuminated by red light) for 1 h before being placed into end-chamber A. Fifteen seconds later, if the rat was facing the door to the middle chamber B, the door was lifted. Videorecording continued for 5 min during free exploration of all 3 chambers. This procedure was performed twice daily, with the probe heights on each trial indicated in each figure. The first three 5-minute trials were performed with 0-mm probe heights, to provide limited familiarization with the MCS apparatus in the absence of painful stimulation. The rats were acclimated to the testing room for 1 hour before each test. In addition, the rats had been habituated previously to the tester and testing room (outside of the MCS) for 6-7 hours over 2 days. After each test in the MCS, the device was sprayed and wiped with Rescue^TM^ disinfectant cleaner (Covetrus). All trials were captured at 1080p, 120 fps using a Sony FDR-AX43 UHD 4K Handycam^®^ camcorder. During the first crossing, the crossing latency was defined as the time from lifting the door between chambers A and B until all four paws were placed in chamber C. Each subsequent crossing was counted when two or more paws and the head of the rat entered the opposite end-chamber, A or C ^54^.

#### Protections against unconscious experimenter bias

Animals were randomly assigned to experimental and control (naïve) groups. All control animals received handling, acclimation, and testing identical to the experimental animals. Guarding behavior was tested by a blinded observer, although during the 1-day post-injury tests the wound was visible to the tester. It was unnecessary to conduct the mechanical conflict (MC) tests blinded for the following reasons: 1) each crossing of the middle chamber is an all-or-none event that is impossible to miss, making counts of the number of crossings extremely reliable, 2) no explicit test stimuli were delivered by the tester, eliminating the possibility of unconscious bias in test stimulus delivery, 3) after initiating the trial, the tester remained at least 6 feet from the MCS device, receiving no apparent attention from animals, all of which had been habituated to the tester, and 4) all trials were videotaped. Nevertheless, a subset of MC tests was conducted by a blinded tester, and some tests were scored from the video records by a blinded observer to confirm unblinded counts of crossings. We have never found any differences in the number of crossings counted by blinded and unblinded observers.

### 2.7. Data analysis

Statistical analyses of raw electrophysiological data and FIBSI output were performed using Prism (GraphPad Software, La Jolla, CA). Summary data are presented as medians, means ± SEM, or percentage of neurons sampled. Datasets were tested for normality with the Shapiro– Wilk test. The behavioral data were normally distributed and tested with two-way ANOVA followed by Bonferroni multiple comparisons tests. The significance levels indicated by asterisks for the behavioral data presented in the figures are *, *P* < 0.05; **, *P* < 0.01; ***, *P* < 0.001; **** *P* < 0.0001. Most of the electrophysiological data were not normally distributed, so in every case we tested the hypothesis that multiple groups differed from the values of a naïve control group using nonparametric Fisher’s exact tests (categorical variables) or Mann-Whitney U tests. To adjust for multiple comparisons, the significance values were lowered appropriately with Bonferroni corrections. All of these tests involved comparisons of a naïve group to 3 other groups, so significance levels indicated by asterisks in the figures are *, *P* < 0.0167; **, *P* < 0.0033; ***, *P* < 0.0003. To indicate possible trends and allow readers to weigh for themselves the risks of type 1 versus type 2 errors in interpretation of the results, we also report any 2-tailed *P* values < 0.10 that did not meet the criteria for significance after Bonferroni corrections.

## 3. Results

### 3.1. Unilateral plantar incision induces bilateral latent hyperactivity in dissociated DRG neurons that lasts at least two weeks

Given the in vivo and in vitro evidence for nociceptor somata being a significant locus for generating SA linked to other forms of pain in rats ^10, 84^ and humans ^51, 52^, we began by asking whether plantar incision enhances the generation of SA in the somata of neurons dissociated from L4-L6 DRGs 1, 7, or 14 days later. Because dissociation removes most of the tonic in vivo neuromodulatory influences that might normally depolarize nociceptor somata under conditions such as bodily injury, we also asked whether plantar incision would enhance the generation of OA during prolonged artificial depolarization to a membrane potential, −45 mV, which may occur naturally under inflammatory conditions and is usually subthreshold for AP generation ^9, 11, 29, 47,53^. On the basis of the reported behavioral and neural effects of the unilateral plantar incision model ^4, 15, 35, 83, 85, 86^, we hypothesized that hyperactivity (SA or latent OA requiring added depolarization to reach AP threshold) in small DRG neurons would last up to 1 week and be restricted to neurons ipsilateral to the injury. Examples of SA and OA during 10-second segments from 30-second recordings at either RMP (SA) or a holding potential of −45 mV (OA) are shown in **Figure 1A**, which also show the DSFs (see following sections) that trigger APs during SA and OA. Whereas none of the neurons sampled from uninjured “naïve” rats exhibited SA at RMP, some of the neurons sampled both ipsilateral and contralateral to the incision site exhibited SA at each time point tested (1, 7, and 14 days post-injury). The proportion of neurons exhibiting SA at RMP was significantly greater than that of naïve controls at every time point tested post-injury (**Fig. 1B)**. The proportion of neurons exhibiting OA at −45 mV (revealing latent hyperactivity) was also significantly increased ipsilaterally compared with naïve controls at each time point (**Fig. 1C**). In contrast to our initial prediction, significant enhancement of the incidence of neurons exhibiting SA and/or OA was also found in contralateral DRG neurons for at least 14 days post-injury (**Fig. 1B, 1C)**. For both SA and OA ipsilateral and contralateral to the incision, the sustained discharge frequencies were low, within the same range (0.02-2 Hz in nearly all cases), as previously described for tonic ongoing discharge (observed over periods of ≥ 30 seconds) in probable nociceptors after isolation from rats, mice, or humans with various neuropathic conditions ^8–11, 42, 52, 53, 74^. These findings show that incisions penetrating skin and muscle induce low-frequency SA and OA in DRG neurons that last longer (for at least 2 weeks) and is more widespread (including contralateral DRG neurons) than predicted from previously reported effects of unilateral plantar incision.

### 3.2. Hyperactivity after unilateral plantar incision is associated with multiple hyperexcitable alterations in dissociated DRG neurons

What are electrophysiological alterations that underlie the somal hyperactivity that persists in dissociated DRG neurons for 1 to 2 weeks after unilateral plantar incision? In terms of membrane potential, tonic SA or OA can be produced by one or more of three alterations in an isolated neuronal soma: sustained depolarization of RMP up to AP threshold, hyperpolarization (reduction) of AP threshold down to RMP, or enhancement of DSFs that transiently bridge the gap between RMP and AP threshold ^53^. Ipsilateral to the incision site, no significant change in RMP was found, although a weak trend for depolarized RMP was present 7 days post-injury (**Fig. 2A**). In this and all other figures, we indicate trends (potentially deserving further investigation) by reporting *P* values <0.10, including values <0.05 that are not statistically significant after Bonferroni corrections for multiple comparisons. Significant hyperpolarization of AP threshold was found 7 days post-injury, along with a trend in that direction 14 days post-injury (**Fig. 2B**). More consistent effects were found on DSFs measured at a holding potential of −45 mV, which were significantly enhanced 1, 7, and 14 days post-injury (**Figs. 1A, 2C**). In addition, the fraction of neurons that had any large DSFs (≥ 5 mV) at a holding potential of −45 mV was significantly increased 7 days post-injury (**Fig. 2D**). Input resistance, which has a major influence on DSF amplitude ^74^, was measured with 0.5-second hyperpolarizing pulses and showed no significant differences from naïve values, although there was a trend for it to increase 7 days post-injury (**Fig. 2E**). There also were no significant differences in input resistance measured with subthreshold 2-second depolarizing pulses (data not shown). Rheobase measured with the 2-second depolarizing current pulses was significantly decreased 1, 7, and 14 days post-injury (**Fig. 2F**).

**Figure 2.**
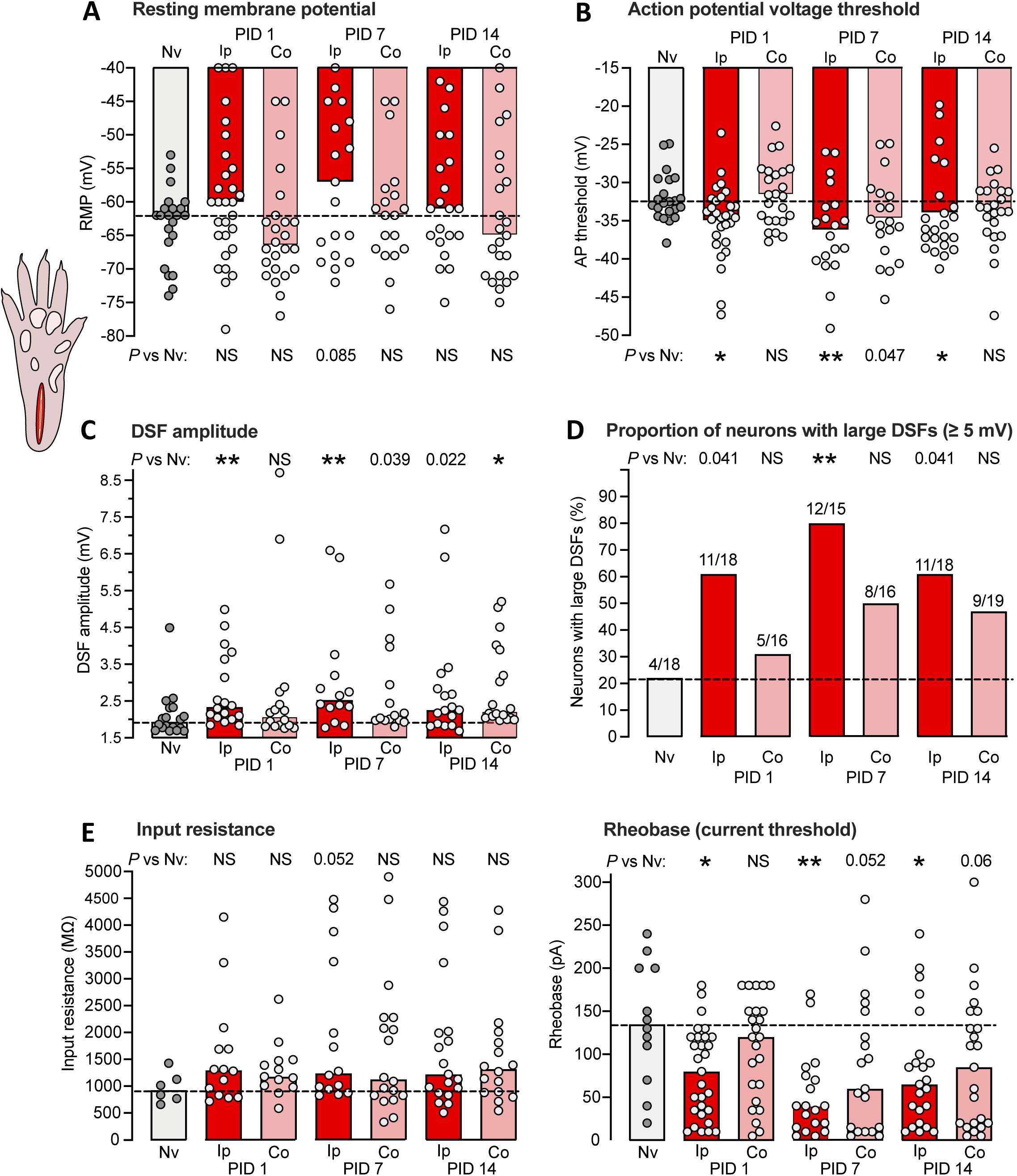
Hyperexcitable alterations found in dissociated DRG neuron somata up to 2 weeks after unilateral plantar incision in vivo. (A) Lack of significant alterations in RMP. In all panels, bars represent medians or proportions; dashed line is the value for the naïve group. Points in a panel represent values for individual neurons. Comparisons by Mann-Whitney U tests with Bonferroni corrections were made for each time point versus naïve group performed separately for the ipsilateral and contralateral groups. Here and in all figures, any numeric *P* values <0.10 are reported to document possible trends. (B) Alterations in AP voltage threshold. Separate comparisons for ipsilateral and contralateral groups were made at each time point versus naïve group by Mann-Whitney U tests with Bonferroni corrections. Note that a numeric *P* value <0.05 reported in the figure means that the comparison did not meet statistical significance after Bonferroni correction. (C) Alterations in DSF amplitude, with separate comparisons made for ipsilateral and contralateral groups at each time point versus naïve group by Mann-Whitney U tests with Bonferroni corrections. (D) Alterations in the proportion of DRG neurons exhibiting large DSFs (≥ 5 mV). Separate comparisons were made for ipsilateral and contralateral groups at each time point versus naïve group by Fisher’s exact tests. (E) Lack of alterations in input resistance assessed by Mann-Whitney U tests with Bonferroni corrections. (F) Alterations in rheobase. Separate comparisons for ipsilateral and contralateral groups at each time point versus naïve group by Mann-Whitney U tests with Bonferroni corrections. AP, action potential; Co, contralateral to injury; DSF, depolarizing spontaneous fluctuation; Ip, ipsilateral; Nv, naïve; NS, not significant; PID, postinjury day.

Dissociated neurons from DRGs contralateral to the incised paw also showed evidence of hyperexcitability after injury. Contralateral DRG neurons exhibited significantly larger DSFs 14 days post-injury (**Fig. 2C**), and trends for effects in the same direction as those in the ipsilateral DRG neurons in AP threshold 7 days post-injury (**Fig. 2B**), in DSF amplitude 7 days post-injury (**Fig. 2C**), and in rheobase 7 and 14 days post-injjury. These results show that unilateral plantar incision can produce bilateral, long-lasting enhancement of DSFs in addition to ipsilateral reduction of AP threshold and rheobase.

### 3.3. Unilateral plantar incision enhances guarding behavior but not avoidance behavior assessed with a mechanical conflict test pitting exploratory activity against pain

The finding of hyperactivity in DRG neurons at unexpected times and locations after unilateral plantar incision raised questions about its relationship to behaviorally expressed pain. Most plantar incision studies have relied on reflexive tests of cutaneous hypersensitivity using withdrawal responses of the injured hindlimb and reductions in either mechanical threshold or the duration of radiant heat stimulation for evoking a response compared to contralateral responses ^60^. Because these segmental nocifensive reflexes can occur independently of affective pain, nocifensive behaviors that depend on emotional processing by the brain are considered better indicators of affective pain experience ^26, 56^. We examined two indicators of affective pain after unilateral paw incision: spontaneous guarding behavior of the incised paw and voluntary avoidance of a noxious substrate revealed with a mechanical conflict (MC) test.

Guarding behavior triggered by plantar incision is sometimes considered an indication of ongoing affective pain and is closely associated with SA in C-fiber and Aδ-fiber nociceptors generated within the injured paw ^83^. Using the same scoring scale, we replicated the finding ^83^ that deep incision significantly increased guarding behavior of the injured hind paw 1 day but not 7 days post-injury when compared to pretest scores (**Fig. 3A**). No guarding was observed in the contralateral hind paw. In contrast to the previous study ^83^, significant guarding behavior was also found 1 day after skin incision alone, perhaps because our rats were not tested 2 hours after injury, and thus had less opportunity to habituate to the testing conditions, or because the rats were tested in the dark phase rather than light phase of their circadian cycle. None of the groups (deep incision, skin incision, or uninjured naive groups) exhibited increased guarding 7 days after injury (**Fig. 3A**). Moreover, a subset of 4 rats examined from each group 14 days after injury showed no differences in mean guarding scores compared to pretest or day 7 scores (not shown). The occurrence of guarding 1 but not 7 days post-injury corresponds to the period reported for mechanical and thermal hypersensitivity after plantar incision ^4, 15, 35, 83, 85, 86^ and to nociceptor SA generated within the hind paw and recorded from teased filaments under general anesthesia in vivo ^4, 5, 83^.

**Figure 3.**
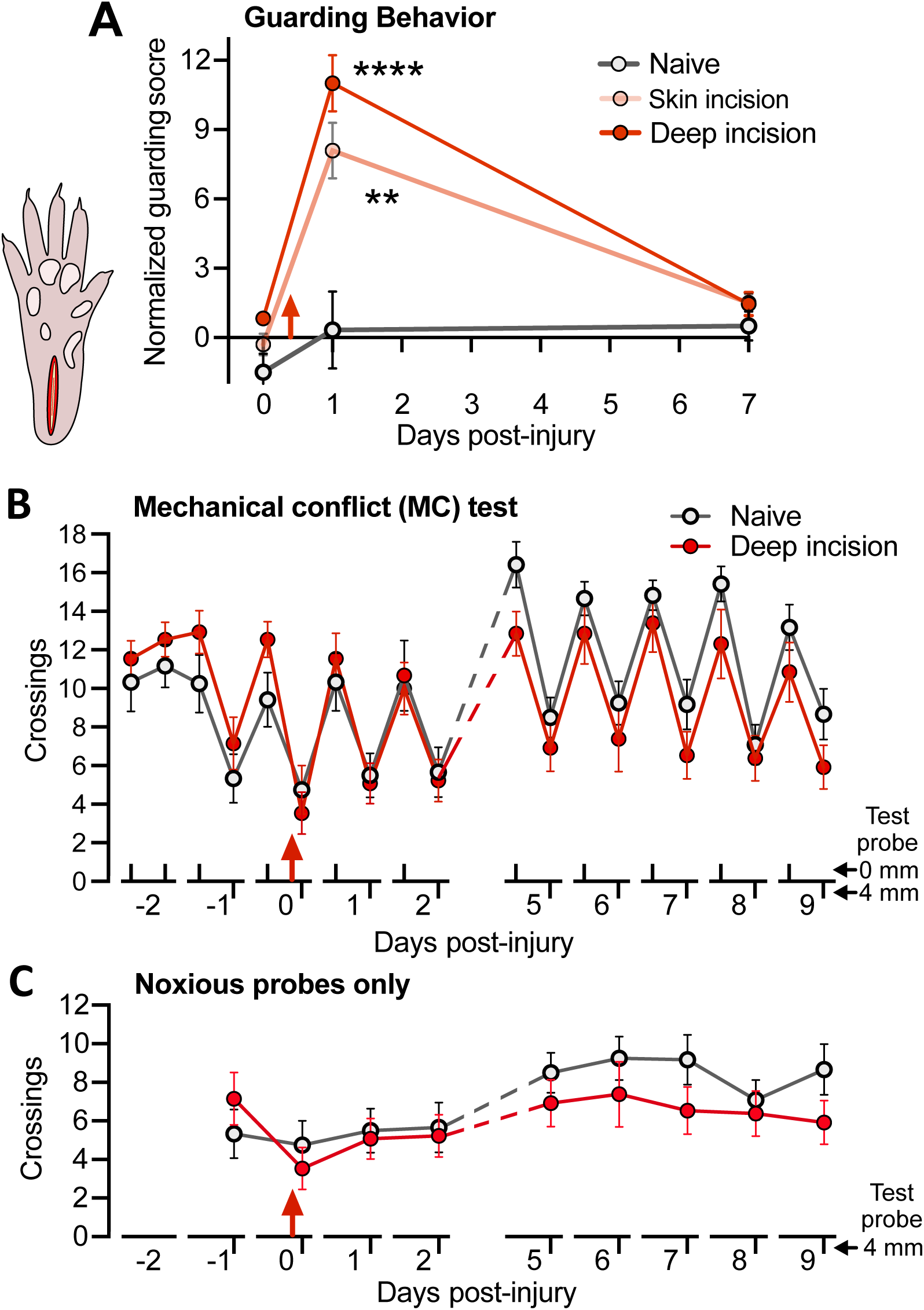
Unilateral plantar incision induces short-lasting guarding behavior but no reduction in crossings of a noxious surface during the operant mechanical conflict (MC) test. (A) Enhancement of mean guarding scores for the injured hindpaw after deep (skin plus muscle) incision or skin incision alone. The guarding score is normalized by subtracting the score of the uninjured hindpaw in each animal. Two-way ANOVA followed by Bonferroni multiple comparison tests: treatment, F(2,79) = 51.5, *P* < 0.0001; test, F(2,79) = 14.0, *P* < 0.0001; asterisks indicate differences compared to pretests on day 0. (B) Lack of effects of unilateral plantar incision on the mean number of crossings of noxious 4-mm probes during 5-minute MC test. Two-way ANOVA followed by Bonferroni tests: test x treatment, F(5,70) = 1.82, *P* = 0.02; test, F(5.74,132) = 20, *P* < 0.0001; treatment, F(1,23) = 4.16, *P* = 0.557. (C) The same data, showing only the crossings of the 4-mm probes. Red arrow indicates the time of incision in each panel.

We used the MC test as our primary indicator of affective pain because it utilizes voluntary behavior (avoidance of noxious probes) to assess increased aversiveness of a noxious stimulus; i.e., mechanical hyperalgesia. Unlike previously reported tests, we omitted the aversive bright light in an end-chamber and relied solely on the rats’ strong exploratory drive to motivate crossing of the middle chamber. Twice-daily 5-minute tests were given, without the probes elevated (0 mm) in the morning and 3 hours later with the probes elevated to 4 mm. The mean number of complete crossings of the middle chamber by the naïve and deep plantar incision groups are plotted across all test days in **Figure 3B**. During the first 3 pretests with the 0-mm probe height, the numbers of crossings were stable and not significantly different between the groups, which then showed parallel decreases in crossings during the single 4-mm probe pretest. Unexpectedly, deep plantar incision produced no statistically significant alterations of crossing numbers during any of the tests, yielding little or no evidence of sensitization or habituation across the 12-day test period. The apparent lack of mechanical hyperalgesia is especially clear when the responses to the 4-mm probes are plotted by themselves (**Fig. 3C**). This surprising result led us to examine more closely our video recordings of the rats’ behavior on the elevated probes. Although the resolution of the recordings was not sufficient for quantitative analysis, two apparent differences in stepping suggested that rats quickly learn to reduce noxious stimulation of the incised heel region of the hind paw when crossing an array of sharp probes. First, longer, higher lifting of the injured hind paw when crossing suggested that the rats shifted weight support to the contralateral paw. Second, the rats often planted the anterior part of the incised hind paw between the probes without lowering the rest of the paw, tiptoeing through the probes and avoiding contact between the sharp probes and the injured heel. These considerations led us to modify the plantar incision model that was used subsequently in order to enable the MC test to reveal potentially long-lasting or rekindled mechanical hyperalgesia.

### 3.4. Bilateral extended plantar incision induces mechanical hyperalgesia manifested as enhanced avoidance behavior that is prolonged by testing for hyperalgesia

Our novel extended plantar incision (EPI) model mitigated limitations of the conventional plantar incision model for indicating affective pain in the MC test. Incision into the plantar flexor digitorum brevis muscle was augmented with a separate, shallower incision across each of the two most anterior footpads (see Methods), and these 3 incisions were made in each of the hind paws. Bilateral EPI makes it much more difficult for a rat to avoid noxious stimulation near an incision site when crossing elevated probes, and it also increases the total amount of tissue damage. Twice-daily 5-minute tests were given, using 1-mm probes in the morning and 4-mm probes 3 hours later. We previously found that, compared to 0-mm probe heights, 1-mm elevations produced no significant reduction in crossings by naïve rats, whereas 4-mm elevations strongly reduced crossings ^54^. Thus, we used reduced voluntary crossing of 1-mm probes as an operant measure of mechanical allodynia and reduced voluntary crossing of 4-mm probes for mechanical hyperalgesia.

The number of crossings of the 1-mm probes was significantly reduced 1 and 5 days after the bilateral EPI procedure compared to naïve controls (**Fig. 4A**). In addition, bilateral EPI significantly reduced crossings of the 4-mm probes 1, 9, and 13 days post-injury, with a strong trend for reduction 5 days post-injury (**Fig. 4B**). These alterations in voluntary crossing of innocuous and noxious surfaces suggest that bilateral EPI induced both mechanical allodynia lasting for close to a week and mechanical hyperalgesia lasting for about 2 weeks.

**Figure 4.**
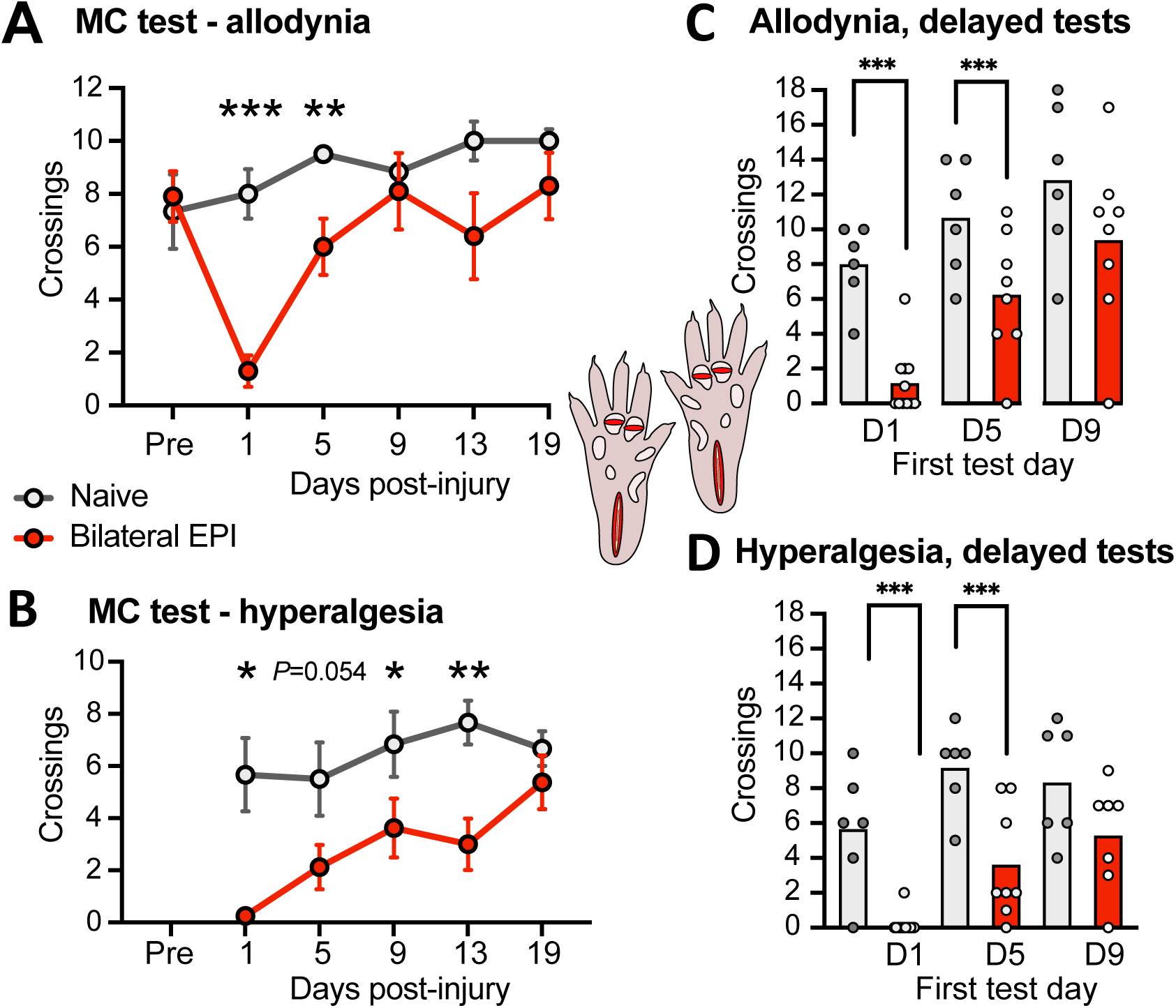
Mechanical allodynia and hyperalgesia after bilateral extended plantar incision (EPI) as revealed by the operant MC test (inset, diagram of the EPI model). (A) Mechanical allodynia indicated by reduced number of crossings of innocuous 1-mm probes. Two-way ANOVA followed by Bonferroni tests: test x treatment, F(5,70) = 4.53, *P* = 0.0012; test, F(2.95, 41.3) = 6.72, *P* = 0.0009; treatment, F(1,14) = 4.16, *P* = 0.060. Asterisks indicate differences between naïve and EPI groups. (B) Mechanical hyperalgesia indicated by reduced number of crossings of noxious 4-mm probes. Two-way ANOVA followed by Bonferroni tests: test x treatment, F(4,56) = 1.81, *P* = 0.139; test, F(2.95, 41.4) = 5.15, *P* = 0.0042; treatment, F(1,14) = 14.75, *P* = 0.0018. Asterisks indicate differences between naïve and EPI groups. (C) Mechanical allodynia when the first test was given 5 but not 9 days post-injury. The hypothesis that allodynia would be present during the first test post-injury was tested with unpaired t-tests for each initial test time (day 5 or day 9) versus the naïve group at the same test time. Results from the first test at day 1 are replotted from panel A for comparison. (D) Mechanical hyperalgesia when the first test was given 5 but not 9 days post-injury. The hypothesis that hyperalgesia would be present during the first test post-injury was tested with unpaired t-tests versus the naïve group at the same initial test time. Results from the first test at day 1 are replotted from panel B for comparison. Averages in all panels are mean values. EPI, extended plantar injury (bilateral); PID, postinjury day.

The allodynia and hyperalgesia observed 5 days or later post-injury might have been influenced by possible nociceptive sensitization from stepping on noxious 4-mm probes during earlier tests. To examine this possibility, we repeated the experiments on separate groups of naïve and EPI rats but delayed the first test until either 5 or 9 days post-injury. Significantly reduced crossings across the 1-mm probes were still found 5 days (and 1 day, replotting the corresponding data from **Fig. 4A**) but not 9 days post-injury (**Fig. 4C**), indicating that bilateral EPI induced allodynia lasted close to 1 week and was independent of previous testing with noxious probes. Significantly reduced crossings across the first test with 4-mm probes were found 1 and 5 days but not 9 days post-injury (**Fig. 4D**). In combination with the significant suppression of crossings at 9 and 13 days post-injury shown in **Figure 4B**, these results suggest that bilateral EPI alone induces mechanical hyperalgesia lasting about 1 week, but repeated testing for hyperalgesia with 4-mm probes adds sensitization that can prolong hyperalgesic behavior for at least 8 more days.

### 3.5. Latent hyperactivity in dissociated DRG neurons persists for 3 weeks after bilateral extended plantar incision

Neurons from bilateral lumbar DRGs (L4-L5) were dissociated 1 day, or 1, 2, 3, or ≥ 4 weeks after bilateral EPI. No differences in the properties of neurons from naïve rats across this extended period were evident, so these neurons were pooled into a single control group. Compared with naïve controls, no significant increases were found in the proportion of neurons exhibiting SA at RMP at any time point (**Fig. 5A**). In contrast, the proportion of neurons exhibiting OA at −45 mV was increased at each time point tested except for 4 weeks or longer post-injury (8% naïve [*N*=13], 79% 1 day [*N*=14], 75% 1 week [*N*=16], 58% 2 weeks [*N*=33], 73% 3 weeks [*N*=15], 10% ≥ 4 weeks [*N*=20]; *P =* 0.0003, 0.0005, 0.0026, 0.0006, respectively, all significant by Fisher’s exact tests versus naïve after Bonferroni corrections). Because the proportion of neurons with OA was approximately the same 1, 2, and 3 weeks post-injury, and these time points occur after the 1-day point when previous plantar incision studies would predict significant primary afferent neuron hyperexcitability, we pooled the data from 1 to 3 weeks post-injury in order to increase statistical power for revealing long-lasting excitability alterations in DRG neurons that are retained after dissociation. The proportion of neurons exhibiting OA at −45 mV 1 day and 1 to 3 weeks post-injury was significantly increased (**Fig. 5B**).

**Figure 5.**
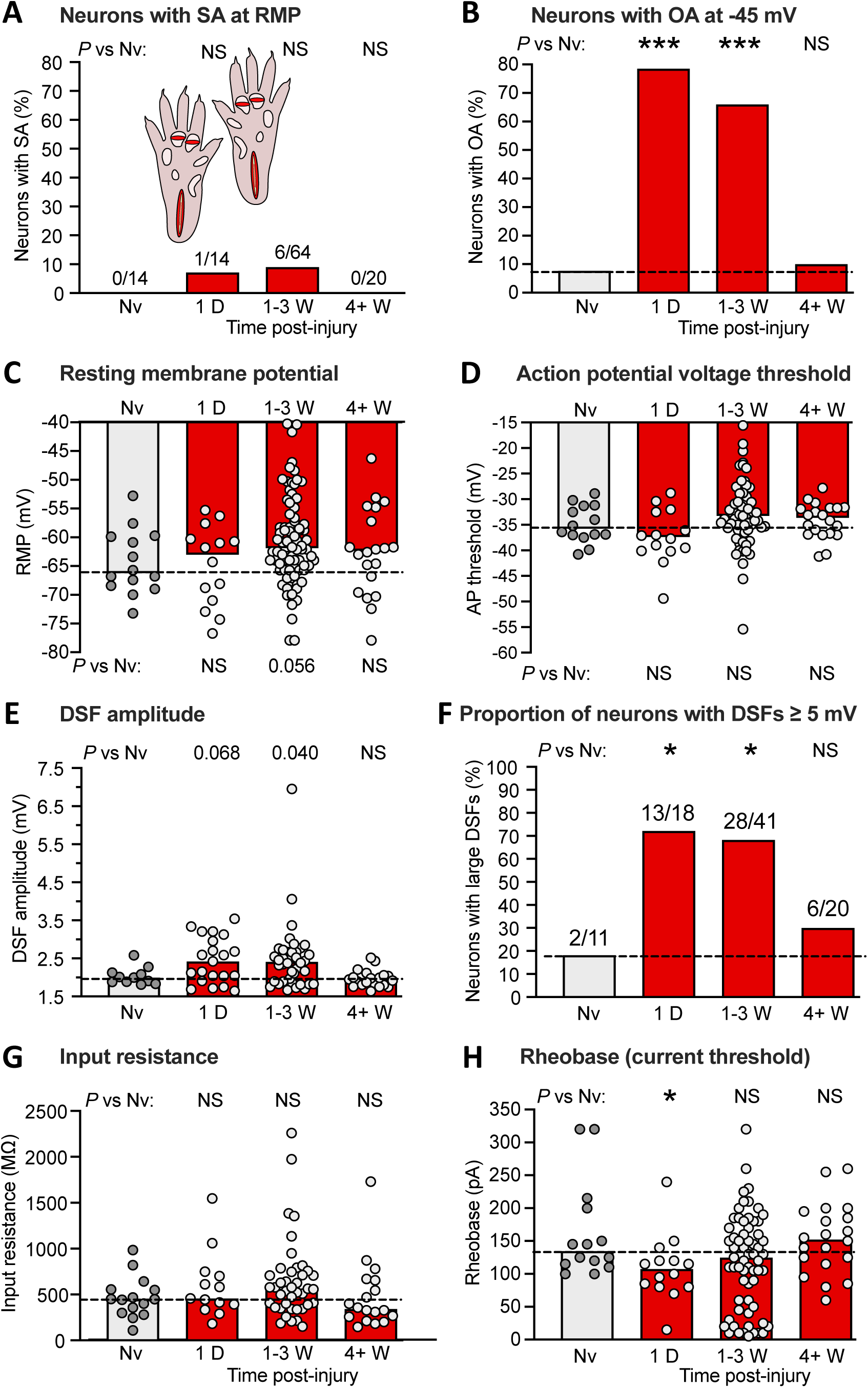
Bilateral extended plantar incision (EPI) induces long-lasting hyperexcitability that is retained after dissociation of DRG neurons. (A) Proportion of neurons exhibiting SA (OA at RMP) when dissociated at the indicated number of postinjury days or weeks following bilateral EPI. Above each bar are the number of neurons exhibiting SA/the number of neurons sampled for that group. Comparisons of EPI with the naïve group at each time period were made with Bonferroni-corrected Fisher’s exact tests. (B) Proportion of the same neurons as in panel A that exhibited OA at −45 mV, assessed with Bonferroni-corrected Fisher’s exact tests. (C) Lack of significant alterations in RMP. Bars and dashed line (naïve group) represent medians. Comparisons by Mann-Whitney U tests with Bonferroni corrections for each time period versus naïve group. (D) Lack of significant alterations in AP voltage threshold as indicated by Mann-Whitney U tests with Bonferroni corrections. (E) Lack of significant alterations in DSF amplitudes (with a possible trend for enhancement). Comparisons were made at each time period versus naïve group with Mann-Whitney U tests with Bonferroni corrections. (F) Alterations in the proportion of DRG neurons exhibiting large DSFs (≥ 5 mV). Comparisons for each time period versus naïve group were made with Fisher’s exact tests. (G) Lack of alterations in input resistance, as assessed wiith Mann-Whitney U tests with Bonferroni corrections. (H) Alterations in rheobase (AP current threshold) assessed for each time period versus naïve group by Mann-Whitney U tests with Bonferroni corrections. AP, action potential; D, day; DSF, depolarizing spontaneous fluctuation; Nv, naïve; NS, not significant; W, weeks.

Compared with naïve control values, no significant differences at any time point after the incisions were found in RMP (**Fig. 5C**) or AP threshold (**Fig. 5D**), although a trend for depolarized RMP was observed 1 to 3 weeks post-injury. Trends for increased DSF amplitude were found 1 day and 1 to 3 weeks post-injury (**Fig. 5E**). The largest effects were found in the proportion of DRG neurons exhibiting large DSFs (≥ 5 mV), with significantly increased incidence of these neurons 1 day and 1 to 3 weeks post-injury (**Fig. 5F**). No significant effects were found on input resistance measured with hyperpolarizing pulses (**Fig. 5G**) or with subthreshold depolarizing pulses (data not shown), while rheobase was significantly reduced 1 day post-injury (**Fig. 5H**).

These results suggest that the most consistent persistent electrophysiological alteration promoting OA at −45 mV after bilateral EPI is an increase in large DSFs, and especially the number of DRG neurons exhibiting large DSFs that may drive widespread, low-frequency tonic activity under conditions that depolarize DRG neurons. Unexpectedly, the number of OA-related excitability properties that were persistently altered was larger after unilateral plantar incision than after bilateral EPI, an observation consistent with the much higher incidence of persistent SA (overt hyperactivity) found in the experiments on unilateral plantar incision (**Fig. 1B, D**) than bilateral EPI (**Fig. 5A**). These differences might be related to differences in the circadian timing between the two sets of experiments; the rats used in the unilateral incision experiments were both injured and later euthanized for DRG collection during the active dark phase of their circadian cycle whereas these manipulations occurred in the inactive light phase in the rats used for bilateral EPI experiments. Other potential differences between sets of experiments could also be at play, such as different personnel handling and testing the rats over the period of this study, and inadvertent differences in DRG neuron isolation, culturing, and recording procedures (e.g., differences between batches of the same reagents, differences in mechanical forces applied by different personnel when triturating cells during dissociation). Small differences in such factors might have substantial effects, especially if subsets of the sampled neurons have stress-detecting functions that make them highly sensitive to various stressors in vivo and/or in vitro. Nevertheless, our results show that the effects of in vivo hindpaw incision are robust enough to consistently enhance OA in subsequently dissociated DRG neurons.

### 3.6. Latent hyperactivity in dissociated DRG neurons after bilateral extended plantar incision occurs in thoracic as well as lumbar DRG neurons

The finding of significant OA at −45 mV in neurons from L4-L6 DRGs contralateral to a unilateral plantar incision (**Fig. 2C, E**) suggested that latent hyperactivity in nociceptors may be a widespread consequence of bodily injury. To explore this possibility further, we examined neurons from more rostral DRGs that innervate dermatomes outside the injured regions on the hind paws. Neurons from bilateral L3-L5 DRGs, T13-L2 DRGs, and T10-T12 DRGs were dissociated in separate groups 1 week after bilateral EPI. No differences in the properties of neurons from naïve rats across these DRG levels were evident, so these neurons were pooled into a single control group. No significant increases were found in the proportion of neurons exhibiting SA at RMP at any of the DRG levels (**Fig. 6A**). The proportion of neurons exhibiting OA at −45 mV was significantly increased in neurons from T13-L2 and T10-T12 DRGs, while a trend was observed in neurons from L3-L5 DRGs (**Fig. 6B**). No significant differences between neurons from any of the DRG levels compared with the naïve group were found in RMP (**Fig. 6C**) or AP threshold (**Fig. 6D**). However, DSF amplitudes (**Fig. 6E**) and the proportion of neurons exhibiting large DSFs (≥ 5 mV) (**Fig. 6F**) were significantly increased in neurons from T13-L2 and T10-T12 DRGs. No significant alterations in neurons from any of the DRG levels were found in input resistance (**Fig. 6G**) or rheobase (**Fig. 6H**).

**Figure 6.**
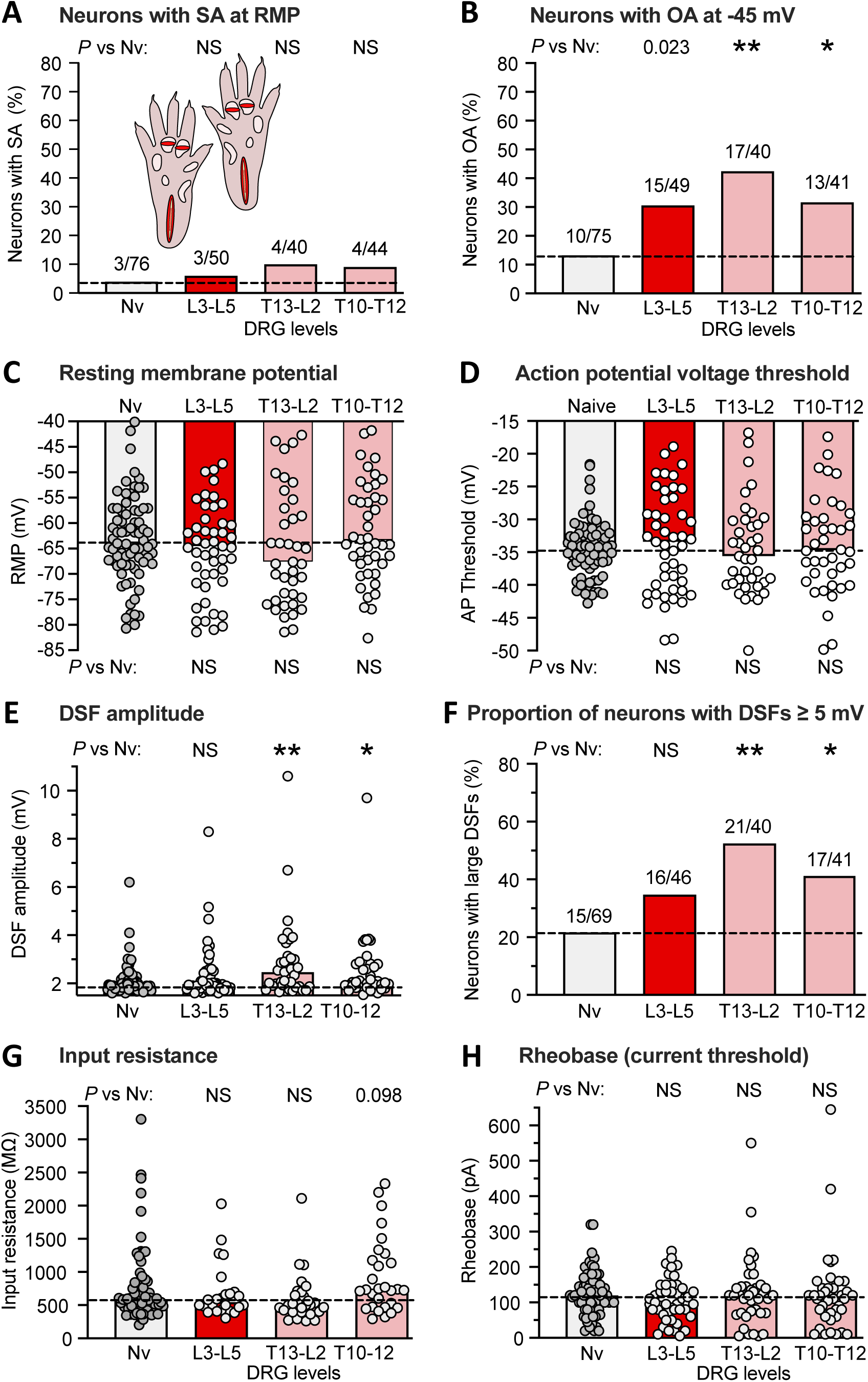
Bilateral extended plantar incision (EPI) induces widespread hyperexcitability of neurons retained after dissociation from thoracic and lumbar DRGs. (A) Proportion of neurons exhibiting SA (OA at RMP) when dissociated from DRGs at the indicated spinal levels. Comparisons of EPI with the naïve group were made at each time period with Bonferroni-corrected Fisher’s exact tests. (B) Proportion of the same neurons as in panel A that exhibited OA at −45 mV assessed with the same statistical tests. (C) Lack of significant alterations in RMP. Bars and dashed line (naïve group) represent medians. Comparisons were made by Mann-Whitney U tests with Bonferroni corrections for each time period versus naïve group. (D) Lack of significant alterations in AP voltage threshold as indicated by Mann-Whitney U tests with Bonferroni corrections. (E) Alterations in DSF amplitudes. Comparisons at each time period versus naïve group were made with Mann-Whitney U tests with Bonferroni corrections at each time period. (F) Alterations in the proportion of DRG neurons exhibiting large DSFs (≥ 5 mV) as assessed with Bonferroni-corrected Fisher’s exact tests. (G) Lack of alterations in input resistance as assessed with Mann-Whitney U tests with Bonferroni corrections. (H) Lack of alterations in rheobase (AP current threshold) as assessed wiith Mann-Whitney U tests with Bonferroni corrections. AP, action potential; DRG, dorsal root ganglion; DSF, depolarizing spontaneous fluctuation; L, lumbar; Nv, naïve; NS, not significant; T, thoracic.

These results confirm that latent hyperactivity, assessed as OA at −45 mV in dissociated DRG neurons, is widespread 1 week after bilateral EPI, occurring in neurons from DRGs that innervate dermatomes distant from the plantar incision sites. The finding of distant latent hyperactivity is striking because in this study the OA and associated excitability alterations in neurons from lumbar DRGs were less pronounced than in the time course study (see **Fig. 5** and Discussion). These results also add to our evidence that enhanced DSFs are the most consistent somal correlate of tissue injury among the excitability properties measured in dissociated DRG neurons. An important implication is that enhancement of DSFs generated within nociceptor somata may be especially important in vivo for latent hyperactivity and possible low-frequency ongoing discharge lasting weeks after bodily injury.

## 4. Discussion

This study revealed latent hyperactivity in DRG neuron somata that is retained for 1 day after dissociation and for weeks after tissue injury occurs. Unexpected findings of somal hyperexcitability in nociceptors isolated from DRGs innervating non-painful dermatomes and persisting after the apparent resolution of pain suggest possible functions of widespread nociceptor hyperactivity in addition to driving ongoing pain.

### 4.1. Nociceptor somata exhibit widespread, long-lasting latent hyperactivity after incision injury

Most neurons examined under our standard experimental conditions are nociceptors, as indicated by their relatively small soma diameters (<30 μm) and previous demonstrations that most are excited by capsaicin and many bind isolectin B4 ^10, 53, 81^. Earlier studies showed that neuropathic insults producing widespread persistent pain, namely SCI ^8–11, 29, 53, 80, 81, 84^ and treatment with the chemotherapeutic agent cisplatin ^42^, induce SA and hyperexcitability that is later expressed in dissociated nociceptor somata. SA and hyperexcitability in dissociated nociceptors have also been linked to neuropathic pain in dermatomes innervated by DRGs or spinal nerves subject to compression by tumors later excised along with the affected DRGs in human cancer patients ^51, 52^. We have now found persistent hyperexcitability of nociceptor somata following peripheral tissue injury.

Three major findings were unexpected. First, somal “memory” of surgical injury in probable nociceptors could be considered unlikely because sensitization and SA generation found in peripheral terminals of primary nociceptors provide a sufficient explanation for driving reflex sensitization and guarding behavior following plantar incision in rats ^4, 5, 60, 83^. Furthermore, transcriptomic ^69, 76^ and proteomic ^61^ analyses of rodent DRGs containing nociceptor somata 1 day after plantar incision have revealed changes in expression of relatively few molecules, which we confirmed with RNA-seq (unpublished data). Very few differentially expressed genes and proteins in these studies directly impact excitability, although one study ^71^ reported upregulation of mRNA and protein for the pain-linked ion channel Nav1.7 after plantar incision. Second, previous studies have found that nociceptor sensitization and SA in nociceptor terminals last less than 1 week ^4, 5, 62, 82, 83^, whereas we find that latent hyperactivity in dissociated nociceptors continues for at least 3 weeks. Third, to our knowledge, all behavioral and electrophysiological alterations reported for rodent plantar incision models have been localized to the incised region. Surprisingly, we found hyperactivity in probable nociceptors having somata in DRGs contralateral and rostral to the DRGs innervating incision sites.

### 4.2. Bilateral extended plantar incision reveals mechanical hyperalgesia manifested in voluntary behavior temporally uncorrelated with nociceptor hyperactivity

An important question concerns the relationship between latent nociceptor hyperactivity and affective pain. Incision studies in rats consistently report that hypersensitivity to weak tactile stimulation and/or hypersensitivity to radiant heat (both utilizing paw withdrawal reflexes) persist for ∼4-10 days and only occur in the injured hind paw ^2, 15, 35, 55, 63, 71, 82, 86^, although additional responses to stimulation of the hypersensitive hind paw may be expressed in other parts of the body ^16, 24^. We chose to examine two behaviors considered better indicators of affective pain than reflexive tests. First, we replicated previous studies ^6, 63, 82, 83^ showing that plantar incision induces guarding behavior that lasts at least 1 day but not 7 days after injury. Although this non-evoked behavior is consistent with transient spontaneous pain, it does not show that an incision-induced state is aversive, which is a defining property of affective pain. To investigate the aversiveness of the central state induced by plantar incision, we used the operant MC test where a rat may voluntarily choose to repeatedly cross an array of sharp probes, with the rat’s drive to explore ^54^ conflicting with pain evoked by the probes. Unilateral plantar incision had no apparent effect on the number of crossings. However, closer inspection of the rats’ behavior suggested that rats minimized noxious stimulation near the incision site by transferring weight to the contralateral hind limb and “tiptoeing” with the injured paw. In contrast, bilateral EPI revealed mechanical allodynia (fewer crossings of 1-mm probes) and hyperalgesia (fewer crossings of 4-mm probes) that lasted at least 5 days but was resolving by 9 days post-injury. The ∼1-week allodynic-hyperalgesic period after bilateral EPI is roughly the same as described previously for sensitization of guarding and withdrawal reflexes after unilateral plantar incision, but less than the ∼3-week period of latent hyperactivity we found in nociceptors after unilateral or bilateral plantar injury. The longer persistence of nociceptor OA demonstrates an important temporal dissociation between latent hyperactivity and pain behavior.

### 4.3. Plantar incision consistently increases the incidence of DRG neurons exhibiting large, low-frequency DSFs

Are there associations between specific electrophysiological properties and the latent hyperactivity revealed by OA at −45 mV? Unlike SCI ^9, 11, 53^, cisplatin treatment ^42^, or neuropathy in humans ^51, 52^, neither unilateral nor bilateral plantar incisions produced all three of the basic excitability alterations that can drive SA in the absence of transient excitatory inputs such as synaptic potentials or sensory generator potentials: depolarized RMP, reduced AP voltage threshold, and enhanced DSFs. We failed to find significant depolarization of RMP in any of our incision experiments. While enhanced OA was consistently found, many of the other excitability properties tested varied across experiments, perhaps because of the sensitivity of nociceptors to varying bodily stressors in vivo or cellular stressors in vitro. AP voltage threshold was significantly reduced after unilateral but not bilateral incision. Rheobase was significantly reduced ipsilaterally at every time point after unilateral incision, but in just one of the two sets of bilateral incision experiments. Most consistently altered was the proportion of DRG neurons exhibiting large DSFs, which was significantly increased ipsilaterally in the unilateral incision experiments and in both sets of bilateral incision experiments. Furthermore, the median amplitude of all DSFs either significantly increased or tended to increase (*P* < 0.05 after Bonferroni corrections) in most groups after unilateral or bilateral incisions. Significant DSF enhancement was also found in DRG neurons contralateral to the unilateral incision site and was significant in DRG neurons from thoracic DRGs innervating dermatomes distant from bilateral plantar incision sites. Enhancement of DSFs was not primarily a consequence of increased input resistance ^74^, which did not significantly increase in any group. Thus, increased incidence of infrequent large DSFs that approach AP threshold ^9, 11, 53, 74^ appear especially important for enhanced low-frequency OA and SA after bodily injury.

### 4.4. What are the functions of latent hyperactivity in nociceptors?

Latent hyperactivity, manifested as tonic, nonaccommodating, low-frequency discharge of nociceptors during modest depolarization delivered by experimental current injection or application of a low concentration of capsaicin, increases after SCI ^11, 53, 80^, cisplatin treatment ^42^, or plantar incision injury. It seems likely to be present after other bodily insults, but tonic OA during prolonged depolarization and the occurrence of large DSFs are rarely examined in studies of DRG neurons. While SA at RMP is likely to be an important contributor to chronic spontaneous pain after SCI ^43, 80, 84^ and perhaps to brief spontaneous pain after plantar incision ^23^, our findings after plantar incision show that an enhanced but latent capacity for hyperactivity remains for weeks after the normalization of ongoing and sensitized pain behavior, and this increased readiness to generate OA occurs in nociceptors from DRGs innervating regions distant from an injury site. While the functions of low-frequency nociceptor hyperactivity subthreshold for evoking pain are unknown, two possibilities are suggested by our findings when combined with other observations.

First, enhanced readiness of nociceptors to generate nonaccommodating OA when tonically depolarized is likely to amplify mechanisms of hyperalgesic priming ^1, 30, 36, 41, 64, 87^ (**Fig. 7**) and to promote ongoing pain under conditions that unmask latent sensitization ^20, 66^. Hyperalgesic priming and latent sensitization may be protective during periods of increased vulnerability after bodily injury and/or infection ^77–79^. In our experiments (**Fig. 4**), injury-induced hyperalgesic priming interacting with further noxious stimulation during repeated crossing of sharp probes in the MC test might have induced sufficient peripheral and central sensitization (promoted by nociceptor OA) to extend hyperalgesia from 5 days to 13 days after plantar injury.

**Figure 7.**
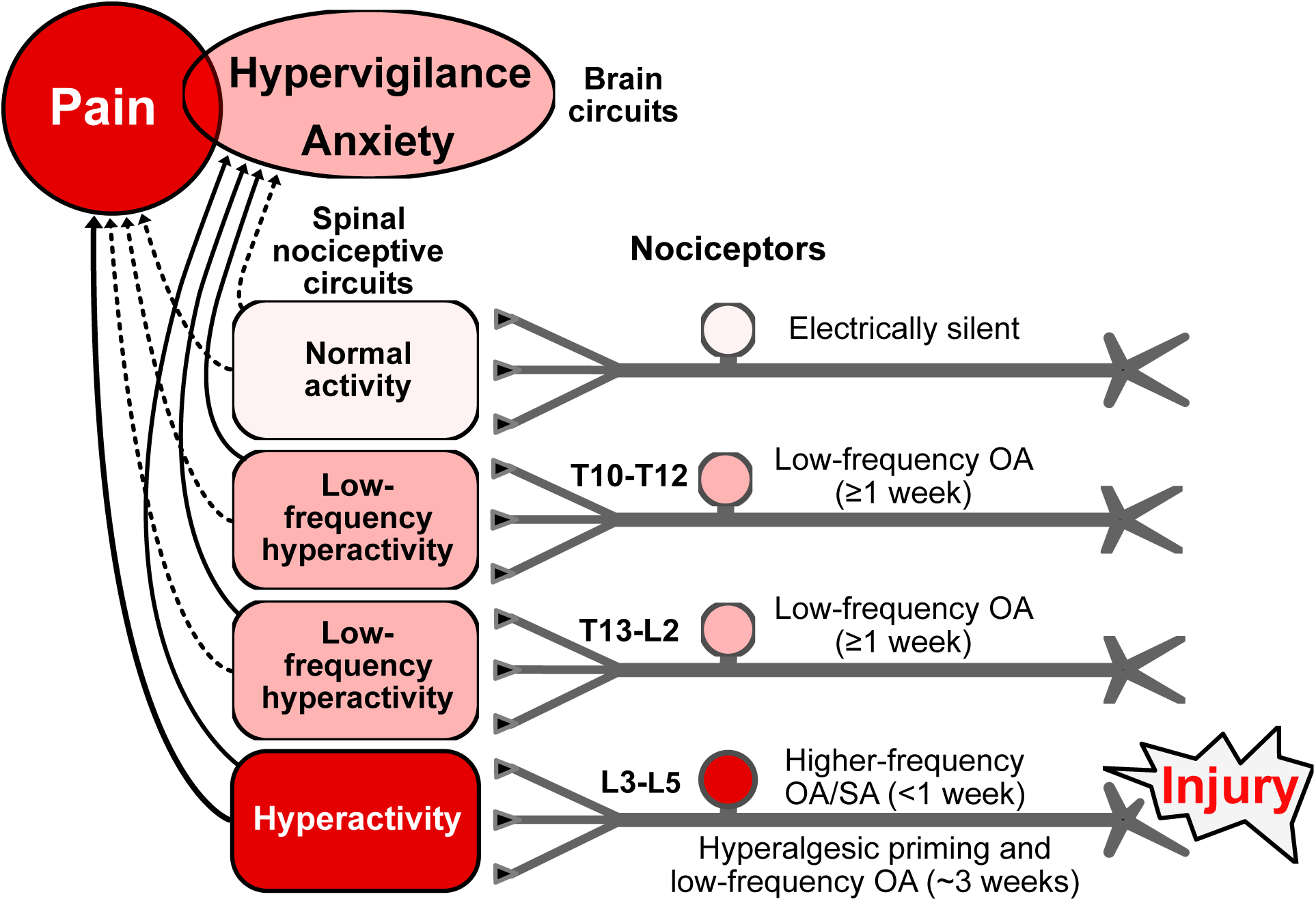
Hypothesis about functions of nociceptor OA based on effects of hindpaw incision injury found to be retained in vitro and on related findings in vivo (see text). Incision induces extrinsic OA stimulated by humoral factors plus intrinsic SA for several days in nociceptors innervating the injured hindpaw (L3-L5 DRGs), with the nociceptor OA activating pathways in the spinal cord that drive pain and/or hypervigilance/anxiety. When ongoing pain ends, lower-frequency OA continues in L3-L5 nociceptors along with hyperalgesic priming for several weeks. Low-frequency OA occurs during the entire ∼3-week period in widespread nociceptors, including those with somata in T13-L2 and T10-T12 DRGs and, if the injury is unilateral, in some contralateral DRGs (not shown). Low-frequency nociceptor OA is proposed to drive general hypervigilance/anxiety more readily than behaviorally significant pain, and to contribute to hyperalgesic priming, especially in the injured region. Higher frequency OA drives ongoing pain as well as anxiety. Red indicates components that drive pain. Pink indicates components with activity subthreshold for driving pain, but which are sufficient to drive hypervigilance/anxiety after hindpaw injury. Solid lines indicate suprathreshold pathways for driving brain-mediated affective states (pain, anxiety, perhaps other negative states); dashed lines are subthreshold. Some electrically silent nociceptors are found in all DRGs after incision injury, but silent nociceptors may be most common in DRGs farthest from the injury. L, lumbar, OA, ongoing activity, T, thoracic.

Second, although nociceptors showing latent hyperactivity in vitro after incision usually require experimental depolarization to reach AP threshold, the same nociceptors in vivo might be tonically depolarized by modulators in the blood and/or CSF to generate low-frequency OA ^77, 78^. An intriguing possibility is that low-frequency OA generated in the somata and terminals of numerous nociceptors may produce diffuse excitation of spinal pathways having a role in addition to stimulation of pain: stimulation of anxiety-related hypervigilance (**Fig. 7**). Anxiety is a common comorbidity of persistent human pain ^18, 34^ and occurs after incision injury in rats ^3, 40, 44^. Anxiety-like hypervigilance that promotes survival after injury has been associated with widespread nociceptor OA in squid ^21, 22^, and analogous hypervigilance was found after nerve injury in mice ^46, 79^. These observations, together with our finding of low-frequency nociceptor OA remaining after the resolution of pain, raise the possibility that spinal pathways driving anxiety-related hypervigilance might be activated by non-painful, low-frequency OA in nociceptors.

## Acknowledgements

This work was supported by National Institute of Neurological Diseases and Stroke Grant NS111521 to E.T. Walters and M.X. Zhu; NS091759 to C.W. Dessauer and E.T. Walters; and the Fondren Chair in Cellular Signaling (E.T. Walters).

We thank Kerry Chu, Ava Porras, Lizeth Robinson, and Michael Y. Zhu for their assistance with some of the experiments, and Jonathan Lee and Gloria Ni for useful improvements to the FIBSI program code.

## References

[1] Araldi D, Bonet IJM, Green PG, Levine JD. Contribution of G-Protein α-Subunits to Analgesia, Hyperalgesia, and Hyperalgesic Priming Induced by Subanalgesic and Analgesic Doses of Fentanyl and Morphine. J Neurosci 2022; 42:1196–210.

[2] Arora V, Morado-Urbina CE, Gwak YS, Parker RA, Kittel CA, Munoz-Islas E, Miguel Jimenez-Andrade J, Romero-Sandoval EA, Eisenach JC, Peters CM. Systemic administration of a β2-adrenergic receptor agonist reduces mechanical allodynia and suppresses the immune response to surgery in a rat model of persistent post-incisional hypersensitivity. Mol Pain 2021; 17:1744806921997206.

[3] Baastrup C, Jensen TS, Finnerup NB. Pregabalin attenuates place escape/avoidance behavior in a rat model of spinal cord injury. Brain Res 2011; 1370:129–35.

[4] Banik RK, Brennan TJ. Spontaneous discharge and increased heat sensitivity of rat C-fiber nociceptors are present in vitro after plantar incision. Pain 2004; 112:204–13.

[5] Banik RK, Brennan TJ. Trpv1 mediates spontaneous firing and heat sensitization of cutaneous primary afferents after plantar incision. Pain 2009; 141:41–51.

[6] Banik RK, Subieta AR, Wu C, Brennan TJ. Increased nerve growth factor after rat plantar incision contributes to guarding behavior and heat hyperalgesia. Pain 2005; 117:68–76.

[7] Baron R, Hans G, Dickenson AH. Peripheral input and its importance for central sensitization. Ann Neurol 2013; 74:630–6.

[8] Bavencoffe A, Li Y, Wu Z, Yang Q, Herrera J, Kennedy EJ, Walters ET, Dessauer CW. Persistent Electrical Activity in Primary Nociceptors after Spinal Cord Injury Is Maintained by Scaffolded Adenylyl Cyclase and Protein Kinase A and Is Associated with Altered Adenylyl Cyclase Regulation. J Neurosci 2016; 36:1660–8.

[9] Bavencoffe AG, Spence EA, Zhu MY, Garza-Carbajal A, Chu KE, Bloom OE, Dessauer CW, Walters ET. Macrophage Migration Inhibitory Factor (MIF) Makes Complex Contributions to Pain-Related Hyperactivity of Nociceptors after Spinal Cord Injury. J Neurosci 2022; 42:5463–80.

[10] Bedi SS, Yang Q, Crook RJ, Du J, Wu Z, Fishman HM, Grill RJ, Carlton SM, Walters ET. Chronic spontaneous activity generated in the somata of primary nociceptors is associated with pain-related behavior after spinal cord injury. J Neurosci 2010; 30:14870–82.

[11] Berkey SC, Herrera JJ, Odem MA, Rahman S, Cheruvu SS, Cheng X, Walters ET, Dessauer CW, Bavencoffe AG. EPAC1 and EPAC2 promote nociceptor hyperactivity associated with chronic pain after spinal cord injury. Neurobiol Pain 2020; 7:100040.

[12] Bohic M, Pattison LA, Jhumka ZA, Rossi H, Thackray JK, Ricci M, Mossazghi N, Foster W, Ogundare S, Twomey CR, Hilton H, Arnold J, Tischfield MA, Yttri EA, St John Smith E, Abdus-Saboor I, Abraira VE. Mapping the neuroethological signatures of pain, analgesia, and recovery in mice. Neuron 2023; 111:2811–2830.e8.

[13] Brazenor GA, Malham GM, Teddy PJ. Can Central Sensitization After Injury Persist as an Autonomous Pain Generator? A Comprehensive Search for Evidence. Pain Med 2022; 23:1283–98.

[14] Brennan TJ. A rat model of postoperative pain. Curr Protoc Pharmacol 2004; Chapter 5:Unit 5.34.

[15] Brennan TJ, Vandermeulen EP, Gebhart GF. Characterization of a rat model of incisional pain. Pain 1996; 64:493–502.

[16] Cameron DM, Brennan TJ, Gebhart GF. Hind paw incision in the rat produces long-lasting colon hypersensitivity. J Pain 2008; 9:246–53.

[17] Cassidy RM, Bavencoffe AG, Lopez ER, Cheruvu SS, Walters ET, Uribe RA, Krachler AM, Odem MA. Frequency-independent biological signal identification (FIBSI): A free program that simplifies intensive analysis of non-stationary time series data. bioRxiv 2020; 2020.05. 29.123042.

[18] Chen T, Wang J, Wang YQ, Chu YX. Current Understanding of the Neural Circuitry in the Comorbidity of Chronic Pain and Anxiety. Neural Plast 2022; 2022:4217593.

[19] Cooper AH, Hedden NS, Corder G, Lamerand SR, Donahue RR, Morales-Medina JC, Selan L, Prasoon P, Taylor BK. Endogenous µ-opioid receptor activity in the lateral and capsular subdivisions of the right central nucleus of the amygdala prevents chronic postoperative pain. J Neurosci Res 2022; 100:48–65.

[20] Corder G, Doolen S, Donahue RR, Winter MK, Jutras BL, He Y, Hu X, Wieskopf JS, Mogil JS, Storm DR, Wang ZJ, McCarson KE, Taylor BK. Constitutive μ-opioid receptor activity leads to long-term endogenous analgesia and dependence. Science 2013; 341:1394–9.

[21] Crook RJ, Dickson K, Hanlon RT, Walters ET. Nociceptive sensitization reduces predation risk. Curr Biol 2014; 24:1121–5.

[22] Crook RJ, Hanlon RT, Walters ET. Squid have nociceptors that display widespread long-term sensitization and spontaneous activity after bodily injury. J Neurosci 2013; 33:10021– 6.

[23] Dalm BD, Reddy CG, Howard MA, Kang S, Brennan TJ. Conditioned place preference and spontaneous dorsal horn neuron activity in chronic constriction injury model in rats. Pain 2015; 156:2562–71.

[24] De Rantere D, Schuster CJ, Reimer JN, Pang DS. The relationship between the Rat Grimace Scale and mechanical hypersensitivity testing in three experimental pain models. Eur J Pain 2016; 20:417–26.

[25] Devor M. Ectopic discharge in Abeta afferents as a source of neuropathic pain. Exp Brain Res 2009; 196:115–28.

[26] Draxler P, Moen A, Galek K, Boghos A, Ramazanova D, Sandkühler J. Spontaneous, Voluntary, and Affective Behaviours in Rat Models of Pathological Pain. Front Pain Res (Lausanne) 2021; 2:672711.

[27] Finnerup NB, Kuner R, Jensen TS. Neuropathic Pain: From Mechanisms to Treatment. Physiol Rev 2021; 101:259–301.

[28] Fuller AM, Bharde S, Sikandar S. The mechanisms and management of persistent postsurgical pain. Front Pain Res (Lausanne) 2023; 4:1154597.

[29] Garza Carbajal A, Bavencoffe A, Walters ET, Dessauer CW. Depolarization-Dependent C-Raf Signaling Promotes Hyperexcitability and Reduces Opioid Sensitivity of Isolated Nociceptors after Spinal Cord Injury. J Neurosci 2020; 40:6522–35.

[30] Garza Carbajal A, Ebersberger A, Thiel A, Ferrari L, Acuna J, Brosig S, Isensee J, Moeller K, Siobal M, Rose-John S, Levine J, Schaible HG, Hucho T. Oncostatin M induces hyperalgesic priming and amplifies signaling of cAMP to ERK by RapGEF2 and PKA. J Neurochem 2021; 157:1821–37.

[31] Goto T, Sapio MR, Maric D, Robinson JM, Domenichiello AF, Saligan LN, Mannes AJ, Iadarola MJ. Longitudinal peripheral tissue RNA-Seq transcriptomic profiling, hyperalgesia, and wound healing in the rat plantar surgical incision model. FASEB J 2021; 35:e21852.

[32] Haroutounian S, Nikolajsen L, Bendtsen TF, Finnerup NB, Kristensen AD, Hasselstrom JB, Jensen TS. Primary afferent input critical for maintaining spontaneous pain in peripheral neuropathy. Pain 2014; 155:1272–9.

[33] Harte SE, Meyers JB, Donahue RR, Taylor BK, Morrow TJ. Mechanical Conflict System: A Novel Operant Method for the Assessment of Nociceptive Behavior. PLoS One 2016; 11:e0150164.

[34] Hooten WM. Chronic Pain and Mental Health Disorders: Shared Neural Mechanisms, Epidemiology, and Treatment. Mayo Clin Proc 2016; 91:955–70.

[35] Kabadi R, Kouya F, Cohen HW, Banik RK. Spontaneous pain-like behaviors are more sensitive to morphine and buprenorphine than mechanically evoked behaviors in a rat model of acute postoperative pain. Anesth Analg 2015; 120:472–8.

[36] Kandasamy R, Price TJ. The pharmacology of nociceptor priming. Handb Exp Pharmacol 2015; 227:15–37.

[37] Kehlet H, Jensen TS, Woolf CJ. Persistent postsurgical pain: risk factors and prevention. Lancet 2006; 367:1618–25.

[38] Khomula EV, Araldi D, Bonet IJM, Levine JD. Opioid-Induced Hyperalgesic Priming in Single Nociceptors. J Neurosci 2021; 41:31–46.

[39] Kleggetveit IP, Namer B, Schmidt R, Helås T, Rückel M, Ørstavik K, Schmelz M, Jørum E. High spontaneous activity of C-nociceptors in painful polyneuropathy. Pain 2012; 153:2040–7.

[40] Kouya F, Iqbal Z, Charen D, Shah M, Banik RK. Evaluation of anxiety-like behaviour in a rat model of acute postoperative pain. Eur J Anaesthesiol 2015; 32:242–7.

[41] Lackovic J, Price TJ, Dussor G. MNK1/2 contributes to periorbital hypersensitivity and hyperalgesic priming in preclinical migraine models. Brain 2023; 146:448–54.

[42] Laumet G, Bavencoffe A, Edralin JD, Huo XJ, Walters ET, Dantzer R, Heijnen CJ, Kavelaars A. Interleukin-10 resolves pain hypersensitivity induced by cisplatin by reversing sensory neuron hyperexcitability. Pain 2020; 161:2344–52.

[43] Lauzadis J, Liu H, Lu Y, Rebecchi MJ, Kaczocha M, Puopolo M. Contribution of T-Type Calcium Channels to Spinal Cord Injury-Induced Hyperexcitability of Nociceptors. J Neurosci 2020; 40:7229–40.

[44] Li CQ, Zhang JW, Dai RP, Wang J, Luo XG, Zhou XF. Surgical incision induces anxiety-like behavior and amygdala sensitization: effects of morphine and gabapentin. Pain Res Treat 2010; 2010:705874.

[45] Li Y, Tatsui CE, Rhines LD, North RY, Harrison DS, Cassidy RM, Johansson CA, Kosturakis AK, Edwards DD, Zhang H, Dougherty PM. Dorsal root ganglion neurons become hyperexcitable and increase expression of voltage-gated T-type calcium channels (Cav3.2) in paclitaxel-induced peripheral neuropathy. Pain 2017; 158:417–29.

[46] Lister KC, Bouchard SM, Markova T, Aternali A, Denecli P, Pimentel SD, Majeed M, Austin JS, de C Williams AC, Mogil JS. Chronic pain produces hypervigilance to predator odor in mice. Curr Biol 2020; 30:R866–7.

[47] Lopez ER, Carbajal AG, Tian JB, Bavencoffe A, Zhu MX, Dessauer CW, Walters ET. Serotonin enhances depolarizing spontaneous fluctuations, excitability, and ongoing activity in isolated rat DRG neurons via 5-HT_4_ receptors and cAMP-dependent mechanisms. Neuropharmacology 2021; 184:108408.

[48] Marvizon JC, Walwyn W, Minasyan A, Chen W, Taylor BK. Latent sensitization: a model for stress-sensitive chronic pain. Curr Protoc Neurosci 2015; 71:9.50.1–14.

[49] Meacham K, Shepherd A, Mohapatra DP, Haroutounian S. Neuropathic Pain: Central vs. Peripheral Mechanisms. Curr Pain Headache Rep 2017; 21:28.

[50] Moy JK, Khoutorsky A, Asiedu MN, Dussor G, Price TJ. eIF4E Phosphorylation Influences Bdnf mRNA Translation in Mouse Dorsal Root Ganglion Neurons. Front Cell Neurosci 2018; 12:29.

[51] North RY, Li Y, Ray P, Rhines LD, Tatsui CE, Rao G, Johansson CA, Zhang H, Kim YH, Zhang B, Dussor G, Kim TH, Price TJ, Dougherty PM. Electrophysiological and transcriptomic correlates of neuropathic pain in human dorsal root ganglion neurons. Brain 2019; 142:1215–26.

[52] North RY, Odem MA, Li Y, Tatsui CE, Cassidy RM, Dougherty PM, Walters ET. Electrophysiological Alterations Driving Pain-Associated Spontaneous Activity in Human Sensory Neuron Somata Parallel Alterations Described in Spontaneously Active Rodent Nociceptors. J Pain 2022; 23:1343–57.

[53] Odem MA, Bavencoffe AG, Cassidy RM, Lopez ER, Tian J, Dessauer CW, Walters ET. Isolated nociceptors reveal multiple specializations for generating irregular ongoing activity associated with ongoing pain. Pain 2018; 159:2347–62.

[54] Odem MA, Lacagnina MJ, Katzen SL, Li J, Spence EA, Grace PM, Walters ET. Sham surgeries for central and peripheral neural injuries persistently enhance pain-avoidance behavior as revealed by an operant conflict test. Pain 2019; 160:2440–55.

[55] Ohta J, Suto T, Kato D, Hiroki T, Obata H, Saito S. Loss of endogenous analgesia leads to delayed recovery from incisional pain in a rat model of chronic neuropathic pain. Brain Res 2020; 1727:146568.

[56] Pahng AR, Edwards S. Measuring Pain Avoidance-Like Behavior in Drug-Dependent Rats. Curr Protoc Neurosci 2018; 85:e53.

[57] Paige C, Plasencia-Fernandez I, Kume M, Papalampropoulou-Tsiridou M, Lorenzo LE, David ET, He L, Mejia GL, Driskill C, Ferrini F, Feldhaus AL, Garcia-Martinez LF, Akopian AN, De Koninck Y, Dussor G, Price TJ. A Female-Specific Role for Calcitonin Gene-Related Peptide (CGRP) in Rodent Pain Models. J Neurosci 2022; 42:1930–44.

[58] Parraga JP, Castellanos A. A Manifesto in Defense of Pain Complexity: A Critical Review of Essential Insights in Pain Neuroscience. Journal of Clinical Medicine 2023; 12:7080.

[59] Pierik JG, IJzerman MJ, Gaakeer MI, Vollenbroek-Hutten MM, van Vugt AB, Doggen CJ. Incidence and prognostic factors of chronic pain after isolated musculoskeletal extremity injury. Eur J Pain 2016; 20:711–22.

[60] Pogatzki-Zahn E, Segelcke D, Zahn P. Mechanisms of acute and chronic pain after surgery: update from findings in experimental animal models. Curr Opin Anaesthesiol 2018; 31:575–85.

[61] Pogatzki-Zahn EM, Gomez-Varela D, Erdmann G, Kaschube K, Segelcke D, Schmidt M. A proteome signature for acute incisional pain in dorsal root ganglia of mice. Pain 2021; 162:2070–86.

[62] Pogatzki EM, Gebhart GF, Brennan TJ. Characterization of Adelta- and C-fibers innervating the plantar rat hindpaw one day after an incision. J Neurophysiol 2002; 87:721–31.

[63] Raithel SJ, Sapio MR, LaPaglia DM, Iadarola MJ, Mannes AJ. Transcriptional Changes in Dorsal Spinal Cord Persist after Surgical Incision Despite Preemptive Analgesia with Peripheral Resiniferatoxin. Anesthesiology 2018; 128:620–35.

[64] Reichling DB, Levine JD. Critical role of nociceptor plasticity in chronic pain. Trends Neurosci 2009; 32:611–8.

[65] Romero A, Romero-Alejo E, Vasconcelos N, Puig MM. Glial cell activation in the spinal cord and dorsal root ganglia induced by surgery in mice. Eur J Pharmacol 2013; 702:126– 34.

[66] Severino A, Chen W, Hakimian JK, Kieffer BL, Gaveriaux-Ruff C, Walwyn W, Marvizón JCG. Mu-opioid receptors in nociceptive afferents produce a sustained suppression of hyperalgesia in chronic pain. Pain 2018; 159:1607–20.

[67] Shepherd AJ, Mohapatra DP. Pharmacological validation of voluntary gait and mechanical sensitivity assays associated with inflammatory and neuropathic pain in mice. Neuropharmacology 2018; 130:18–29.

[68] Shin SM, Cai Y, Itson-Zoske B, Qiu C, Hao X, Xiang H, Hogan QH, Yu H. Enhanced T-type calcium channel 3.2 activity in sensory neurons contributes to neuropathic-like pain of monosodium iodoacetate-induced knee osteoarthritis. Mol Pain 2020; 16:1744806920963807.

[69] Spofford CM, Brennan TJ. Gene expression in skin, muscle, and dorsal root ganglion after plantar incision in the rat. Anesthesiology 2012; 117:161–72.

[70] Study RE, Kral MG. Spontaneous action potential activity in isolated dorsal root ganglion neurons from rats with a painful neuropathy. Pain 1996; 65:235–42.

[71] Sun J, Li N, Duan G, Liu Y, Guo S, Wang C, Zhu C, Zhang X. Increased Na_v_1.7 expression in the dorsal root ganglion contributes to pain hypersensitivity after plantar incision in rats. Mol Pain 2018; 14:1744806918782323.

[72] Suzuki T, Shimizu T, Karnup S, Shimizu N, Ni J, de Groat WC, Yoshimura N. Therapeutic effects of p38 mitogen-activated protein kinase inhibition on hyperexcitability of capsaicin sensitive bladder afferent neurons in mice with spinal cord injury. Life Sci 2023; 325:121738.

[73] Tappe-Theodor A, King T, Morgan MM. Pros and Cons of Clinically Relevant Methods to Assess Pain in Rodents. Neurosci Biobehav Rev 2019; 100:335–43.

[74] Tian J, Bavencoffe AG, Zhu MX, Walters ET. Readiness of nociceptor cell bodies to generate spontaneous activity results from background activity of diverse ion channels and high input resistance. Pain 2023;

[75] Tode J, Kirillova-Woytke I, Rausch VH, Baron R, Jänig W. Mechano- and thermosensitivity of injured muscle afferents 20 to 80 days after nerve injury. J Neurophysiol 2018; 119:1889–901.

[76] Tran PV, Johns ME, McAdams B, Abrahante JE, Simone DA, Banik RK. Global transcriptome analysis of rat dorsal root ganglia to identify molecular pathways involved in incisional pain. Mol Pain 2020; 16:1744806920956480.

[77] Walters ET. Nociceptors as chronic drivers of pain and hyperreflexia after spinal cord injury: an adaptive-maladaptive hyperfunctional state hypothesis. Front Physiol 2012; 3:309.

[78] Walters ET. Adaptive mechanisms driving maladaptive pain: how chronic ongoing activity in primary nociceptors can enhance evolutionary fitness after severe injury. Philos Trans R Soc Lond B Biol Sci 2019; 374:20190277.

[79] Walters ET, Crook RJ, Neely GG, Price TJ, Smith ESJ. Persistent nociceptor hyperactivity as a painful evolutionary adaptation. Trends Neurosci 2023; 46:211–27.

[80] Wu Z, Li L, Xie F, Du J, Zuo Y, Frost JA, Carlton SM, Walters ET, Yang Q. Activation of KCNQ Channels Suppresses Spontaneous Activity in Dorsal Root Ganglion Neurons and Reduces Chronic Pain after Spinal Cord Injury. J Neurotrauma 2017; 34:1260–70.

[81] Wu Z, Yang Q, Crook RJ, O’Neil RG, Walters ET. TRPV1 channels make major contributions to behavioral hypersensitivity and spontaneous activity in nociceptors after spinal cord injury. Pain 2013; 154:2130–41.

[82] Xu J, Brennan TJ. Comparison of skin incision vs. skin plus deep tissue incision on ongoing pain and spontaneous activity in dorsal horn neurons. Pain 2009; 144:329–39.

[83] Xu J, Brennan TJ. Guarding pain and spontaneous activity of nociceptors after skin versus skin plus deep tissue incision. Anesthesiology 2010; 112:153–64.

[84] Yang Q, Wu Z, Hadden JK, Odem MA, Zuo Y, Crook RJ, Frost JA, Walters ET. Persistent pain after spinal cord injury is maintained by primary afferent activity. J Neurosci 2014; 34:10765–9.

[85] Yoshiyama Y, Sugiyama Y, Ishida K, Fuseya S, Tanaka S, Kawamata M. Plantar incision with severe muscle injury can be a cause of long-lasting postsurgical pain in the skin. Life Sci 2021; 275:119389.

[86] Zahn PK, Brennan TJ. Primary and secondary hyperalgesia in a rat model for human postoperative pain. Anesthesiology 1999; 90:863–72.

[87] Zhang Y, Jeske NA. A-kinase anchoring protein 79/150 coordinates α-amino-3-hydroxy-5-methyl-4-isoxazolepropionic acid receptor sensitization in sensory neurons. Mol Pain 2023; 19:17448069231222406.

